# Distinct neurocognitive bases for social trait judgments of faces in autism spectrum disorder

**DOI:** 10.1101/2021.11.18.469134

**Authors:** Hongbo Yu, Runnan Cao, Chujun Lin, Shuo Wang

## Abstract

Autism spectrum disorder (ASD) is characterized by difficulties in social processes, interactions, and communication. Yet, the neurocognitive bases underlying these difficulties are unclear. Here, we triangulated the ‘trans-diagnostic’ approach to personality, social trait judgments of faces, and neurophysiology to investigate (1) the relative position of autistic traits in a comprehensive social-affective personality space and (2) the distinct associations between the social-affective personality dimensions and social trait judgment from faces in individuals with ASD and neurotypical individuals. We collected personality and facial judgment data from a large sample of online participants (*N* = 89 self-identified ASD; *N* = 308 neurotypical controls). Factor analysis with 33 sub-scales of 10 social-affective personality questionnaires identified a 4-dimensional personality space. This analysis revealed that ASD and control participants did not differ significantly along the personality dimensions of empathy and prosociality, antisociality, or social agreeableness. However, the associations between these dimensions and judgments of facial trustworthiness and warmth differed across groups. Neurophysiological data also indicated that ASD and control participants might rely on distinct neuronal representations for judging trustworthiness and warmth from faces. These results suggest that the atypical association between social-affective personality and social trait judgment from faces may contribute to the social and affective difficulties associated with ASD.

## Introduction

Autism spectrum disorder (ASD) is defined by difficulties in social processes, interactions, and communication (APA, 2013). However, there are substantial inter-individual variabilities in terms of both the intensity of the difficulties and the specific aspects of social cognitive and affective functioning that are impaired (Martinez-Murcia et al., 2017; Mottron & Bzdok, 2020). Diagnosing an individual with ASD does not explain the difficulties in their social cognitive and affective functions (e.g., understanding others’ emotion and intention, engaging reciprocally). What psychological mechanisms might underlie the social-affective difficulties manifested in ASD?

To address this important question, various psychological mechanisms have been proposed. Some researchers have found that alexithymia, the difficulty in recognizing and describing one’s own and others’ emotional states (Brewer et al., 2016; Grynberg et al., 2012; Nemiah, 1976), can explain the difficulties with social interactions and emotional reciprocity observed in people with ASD (Bird et al., 2011; Bird & Cook, 2013; Cuve, Castiello, et al., 2021; Cuve, Murphy, et al., 2021). Others have suggested that deficits in empathy (Baron-Cohen & Wheelwright, 2004; McDonald & Messinger, 2012; Rogers et al., 2007; Rueda et al., 2015), the ability to vicariously experience another’s feelings and be concerned about another’s suffering, may underlie the impairments in social interactions that are central to ASD, such as difficulties with emotional engagement (APA, 2013).

Although these previous studies have revealed some correlates of the social-affective difficulties manifested in ASD, to systematically understand the psychological mechanisms underlying them, it is essential to characterize the relative position of autistic traits in a comprehensive social-affective personality space (e.g., empathy, anxiety, prosociality, and antisociality). Extant studies typically focused on only one or two personality measures (e.g., alexithymia, or empathy, as mentioned above), without controlling for other covarying personality constructs. This may lead to an imprecise and incomplete understanding of the personality profile of ASD (see (Nicholson et al., 2018; Shah et al., 2019)). The limited scope of the personality measures examined also contributes to the problem of biased samples in prior studies. For example, some studies recruited participants with ASD and typically developing (TD) control participants that were matched in alexithymia in order to dissociate the contributions of alexithymia and autistic traits to the social-affective difficulties observed in individuals with ASD (e.g., (Bird et al., 2010; Bird & Cook, 2013)). Due to the difference in the baseline prevalence of alexithymia in the TD (5%) and the ASD (50%) populations (Kinnaird et al., 2019) (Kinnaird et al., 2019), the resultant groups were therefore potentially biased and not representative of their respective populations (see (Shah et al., 2019)). Therefore, understanding the relationships of autistic traits to a comprehensive set of social-affective personalities is critical for both ascertaining what social-affective personality dimensions are specifically impaired in autistic individuals and which dimensions are comparable across ASD and TD individuals, and guiding more representative sampling of participants.

For a social-affective personality dimension to have any measurable consequences, it needs to be manifested in some behavioral performance, neural response patterns, or both. In the literature, behavioral consequences of the difficulties associated with autistic traits have been assessed predominantly using emotion recognition tasks with static human facial pictures depicting the so-called ‘basic emotions’ (Adolphs et al., 2001; Pelphrey et al., 2002). Extensive research adopting this approach has shown that people with ASD have pervasive impairments in recognizing facial expressions from static facial pictures (see (Webster et al., 2021) for a review). Such impairments may result from their relatively limited amounts of time spent on, and atypical attention patterns in, viewing human faces (Adolphs et al., 2001; Dawson et al., 2005; Kliemann et al., 2010; Klin et al., 2002; Neumann et al., 2006; Pelphrey et al., 2002; Spezio et al., 2007b, 2007a).

However, social-affective performance is more diverse than recognizing facial expressions from faces (Jack & Schyns, 2017; Todorov et al., 2015). People readily and rapidly make judgments regarding the social traits of a person merely from the appearance of their face (i.e., temporally stable characteristics, such as warmth, trustworthiness, and competence). These judgments have profound consequences for interpersonal interactions and collective decisions in politics and justice (Fiske et al., 2007; C. Lin et al., 2018; Todorov et al., 2015). For example, perceiving a face as more trustworthy is associated with higher financial investments in trust games, regardless of actual trustworthiness (Rezlescu et al., 2012). Perceiving a face as physically attractive leads to the inference of competence and intelligence (Eagly et al., 1991), and the perceived warmth of a face is associated with liking (Wojciszke et al., 2009). In contrast, observers assign harsher sentences to inmates with more Afrocentric features than those with less Afrocentric features (R J R Blair et al., 2004). Although there is a plethora of literature showing abnormal emotion judgment (Webster et al., 2021) and gaze patterns on faces in ASD (Kliemann et al., 2010; Klin et al., 2002; Neumann et al., 2006; Pelphrey et al., 2002), social trait judgment from faces in ASD remains largely unexplored. The few existing studies reported mixed findings. While some studies reveal abnormal social trait judgments regarding the trustworthiness of faces in ASD (Adolphs et al., 2001; Forgeot dArc et al., 2016), others argue that people with ASD have largely normal social trait judgments, including trustworthiness (Latimier et al., 2019; Lindahl, 2017). The discrepancy in the literature has not been resolved given the small number of participants involved.

In this study, we aimed to address two critical questions: (1) where are autistic traits situated in a comprehensive social-affective personality space? and (2) what personality dimensions in this social-affective personality space can account for atypical social trait judgments from faces in individuals with ASD. To address the first question, we adopted a dimensional (or ‘trans-diagnostic’) approach to personality measures, applying factor analysis to 33 sub-scales of various partially overlapping social-affective personality questionnaires (Gillan et al., 2016). For the second question, we used naturalistic face images of celebrities taken in real world contexts that have been well validated for social trait judgment in both ASD and control participants in recent studies (Cao et al., 2020, 2021)), and explained the individual differences in social trait judgment using the resultant factor scores identified in question (1). Among the various social trait judgments identified in these previous studies (Cao et al., 2020, 2021), we were particularly interested in the differences in trustworthiness and warmth judgment between the ASD and the control participants. Both of these traits are critically involved in social approach tendencies (Slepian et al., 2012; Slepian & Bastian, 2017; Todorov, 2008), and are therefore most relevant to the difficulties with social communications and interactions that people with ASD exhibit.

## Materials and Methods

### Participants

We acquired social trait ratings of faces from four groups of participants: (1) 412 participants from the general population (mean age = 26.2 years, s.d. = 6.9; 148 female), (2) 113 participants from the general population with self-identified autism spectrum disorder (ASD) (mean age = 28.9 years, s.d. = 8.4; 59 females), (3) 8 neurosurgical patients (5 female) who had undergone surgery to have electrodes implanted to treat intractable epilepsy, and (4) 16 high-functioning participants with ASD (mean age = 23.2, s.d. = 4.5; 2 females) from our laboratory registry who met the DSM-V and Autism Diagnostic Observation Schedule (ADOS) criteria for ASD. The data collection and some data analysis related to (1) and (2) were preregistered (https://aspredicted.org/CMY_BBC). Participants in (1) and (2) were recruited from an online data collection platform Prolific. One-hundred and four participants from (1) and 24 participants from (1) (2) were excluded due to failures in the attention check questions, as we preregistered. The study was approved by the Institutional Review Board of West Virginia University (WVU). Importantly, because the participants in (2) were self-identified as ASD, the accuracy of which we could not verify, we further recruited (4) as a comparison group. We not only confirmed that the AQ (two-tailed two-sample *t*-test: *t*(99) = 0.88, *p* = 0.38) and SRS (*t*(99) = 0.36, *p* = 0.72) scores of the participants in (2) were comparable to those from the in-lab participants with diagnosed ASD (**Fig. S2a, Fig. S2b**), but also showed that the ratings of the same 500 faces (for details, see **Face judgment task** below) from the in-lab participants with ASD were more similar to those from the online participants with ASD than to the online controls (**Figs. S2c-e**). All results reported here are from groups (1) and (2), which we refer to as “control” and “ASD” hereafter. Results from groups (3) and (4) were reported in the *Supplementary Materials*, which corroborated all findings reported here.

### Self-reported personality questionnaires

Participants completed a battery of personality questionnaires assessing their social-affective traits. They can be roughly classified into four categories: (1) affective deficits, including Social Anxiety (Leary, 1983), Apathy (Ang et al., 2017), Alexithymia (Bagby et al., 1994), and Moral Scrupulosity (Summers & Sinnott-Armstrong, 2019), (2) antisocial traits, including the Dark Factors (Moshagen et al., 2020) and Utilitarianism (Kahane et al., 2018), (3) the Big Five (short version; (Rammstedt & John, 2007)), and (4) other-oriented and empathic traits, including QCAE (Reniers et al., 2011), Perceived Social Support (M. Lin et al., 2019), and Prosocialness (Caprara et al., 2005). Participants also provided demographic information, including their age, sex assignment at birth, highest level of education, and subjective social economic status (SES).

### Face judgment task

We used photos of celebrities from the CelebA dataset (Liu et al 2015). We selected 50 identities with 10 images for each identity, totaling 500 face images. The identities were selected to include both genders and multiple races. The faces were of different angles and gaze directions, with diverse backgrounds and lighting. The faces showed various expressions, with some having accessories such as sunglasses and hats.

Participants were asked to rate the faces on ten social traits using a 7-point Likert scale through an online rating task. The social traits included *warm, critical, competent, practical, feminine, strong, youthful, charismatic, trustworthy, and dominant*. The judgments of these social traits from faces were well validated in a previous study (C. Lin et al., 2021; Oosterhof & Todorov, 2008). We used the same stimuli for neural recordings.

We divided the experiment into 10 modules, with each module containing one face image randomly selected per face identity (totaling 50 face images per module). In each module participants rated each face on 10 social traits (e.g., competence; rated in blocks). We applied the following three exclusion criteria prior to statistical analysis:

1. Trial-wise exclusion: we excluded trials with reaction times shorter than 100 ms or longer than 5000 ms.
2. Block/trait-wise exclusion: we excluded the entire block per participant if more than 30% of the trials were excluded from the block per (1) above, or if there were fewer than 3 different rating values in the block (this suggests that the participant may not have used the rating scale properly).
3. Participant-wise exclusion: we excluded a participant if more than 3 blocks were excluded from the participant per (2) above.

Based on these criteria, less than 5% (4.84%±5.69%) of trials were excluded per participant.

### Single-neuron recordings and neuronal response to faces

We recorded from implanted depth electrodes in the amygdala and hippocampus from patients with pharmacologically intractable epilepsy. Target locations in the amygdala and hippocampus were verified using post-implantation CT. At each site, we recorded from eight 40 μm microwires inserted into a clinical electrode as described previously (Rutishauser et al., 2006, 2013). Details of the neural recording can be found in the *Supplementary Methods*.

### Visualizing networks of personality traits

As an exploratory step towards clarifying the relationships between autistic traits and other social-affective personalities, we carried out a series of network analyses based on participants’ scores on the questionnaire sub-scales. The purpose of this analysis was to better illustrate how different personality measures were correlated, and to visualize the relative position of autistic traits among related social affective personality traits based on the correlation structure. We generated network plots using the *R* function “network_plot” in the package *corrr* (cf. (Petitet et al., 2021)). This analysis relies on multidimensional clustering to estimate the statistical distance between variables. This approach provides an intuitive way of illustrating latent clusters in the correlation matrices, namely subsets of variables that are more strongly, either positively or negatively, correlated with one another. Past research has predominantly focused on the relationship between autistic traits and subsets of social affective personalities separately (e.g., (Bird & Cook, 2013)). Although this line of research has identified both positive or negative associations, we still do not have a comprehensive picture of where autistic traits stand in the comprehensive space of social affective traits. To address this question, we categorized our personality measures (other than the autistic trait measures) into four groups. We then subjected autistic traits on the one hand, and each of these groups of personality measures on the other hand, to network analysis. The four categories are: Affective Deficits (Social Anxiety, Apathy, Alexithymia, and Moral Scrupulosity), Antisocial Traits (the Dark Factors and the Utilitarianism), the Big Five, and Other-oriented and Empathic Traits (QCAE, Perceived Social Support, and Prosocialness). The categorization was based on how similar the questionnaires are in terms of their content and the analysis was for illustration purposes.

### Factor analysis

Some of the questionnaires we used to measure social-affective personality overlap conceptually and statistically, rendering it difficult to estimate the specificity of the association between any one personality measure and social trait judgments. To address this issue, we leveraged the dimension approach to personality traits in computational psychiatry (Gillan et al., 2016). Thirty-three questionnaire sub-scales were included in the exploratory factor analysis. The number of factors was determined based on the Cattell-Nelson-Gorsuch (CNG) test (Gorsuch & Nelson, 1981) implemented in the nFactors package in *R* (Raiche & Magis, 2010). Factor analysis model was estimated using the factanal() function in R, with an oblique rotation (oblimin).

### Representational similarity analysis (RSA)

Dissimilarity matrices (DMs) are symmetric matrices of dissimilarity between all pairs of face identities (Kriegeskorte et al., 2008). For each social trait, we first calculated the consensus ratings of the 500 images by averaging the ratings from the control and ASD group separately. The dissimilarity between each identity pair was then measured using the Pearson correlation across the ratings (z-scored) of the 10 face examples of each identity. Correspondingly, for the neural DM, we averaged the responses (firing rates were normalized to the mean baseline of each neuron) of the same 500 images across individual neurons and measured the neural dissimilarity for each identity pair. In a DM, larger values represent larger dissimilarity of pairs, such that the smallest value possible is the similarity of a condition unto itself (dissimilarity of 0). To compare the pattern similarity between each social trait and neural response, we used the Spearman correlation to calculate the correspondence between the DMs. Spearman correlation was used because it does not assume a linear relationship between variables. We further used a permutation test with 1000 runs to statistically compare the DM correspondence between participants with ASD and controls. In each run, we shuffled the participant labels and calculated the difference in DM correspondence between participant groups. We then compared the observed difference in DM correspondence between participant groups with the permuted null distribution to derive statistical significance.

## Results

All de-identified data and data analysis codes related to the results reported in this paper can be accessed at Open Science Framework upon acceptance. We have reported all measures, conditions, and data exclusions. The data collection of personality and some data analyses related to personality and social trait judgment were preregistered (https://aspredicted.org/CMY_BBC).

### Relative position of autistic traits in a comprehensive social-affective personality network

As Figure 2 shows, autistic traits, as measured by AQ and SRS, appeared separate from anti-social traits (e.g., Dark Factors) and other-oriented and empathic traits (e.g., both emotional and cognitive empathy). This supports the notion that autistic traits are independent of (i.e., ***a***social and ***a***moral) rather than antithetical to (i.e., ***anti***-social or ***im***moral) prosociality and other-oriented tendencies (Jaswal & Akhtar, 2019). In contrast, autistic traits were more closely related to being socially anxious (i.e., social anxiety) and difficulty in describing and identifying one’s own emotions (i.e., alexithymia). Similarly, autistic traits were also more closely related to neuroticism and, conversely, to extraversion. This suggests that autistic traits may operate on the same dimension as social avoidance tendencies.

**Figure 1.**
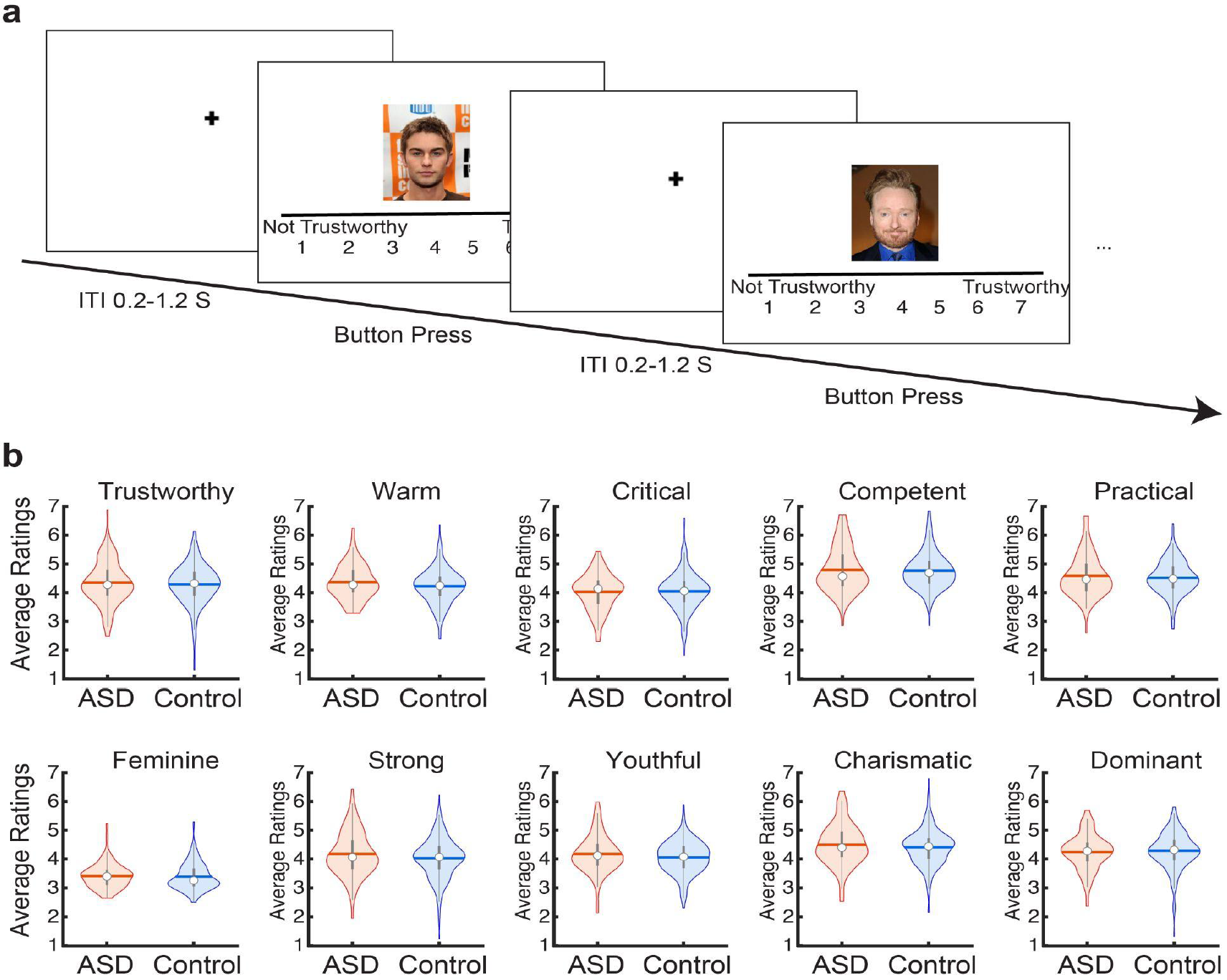
Procedure of the face judgment task **(a)** and behavioral results **(b)**. The mean ratings for the ten social traits were comparable between groups. Violin plots present the median value as the white circle and the interquartile range as the gray vertical bars.

**Figure 2.**
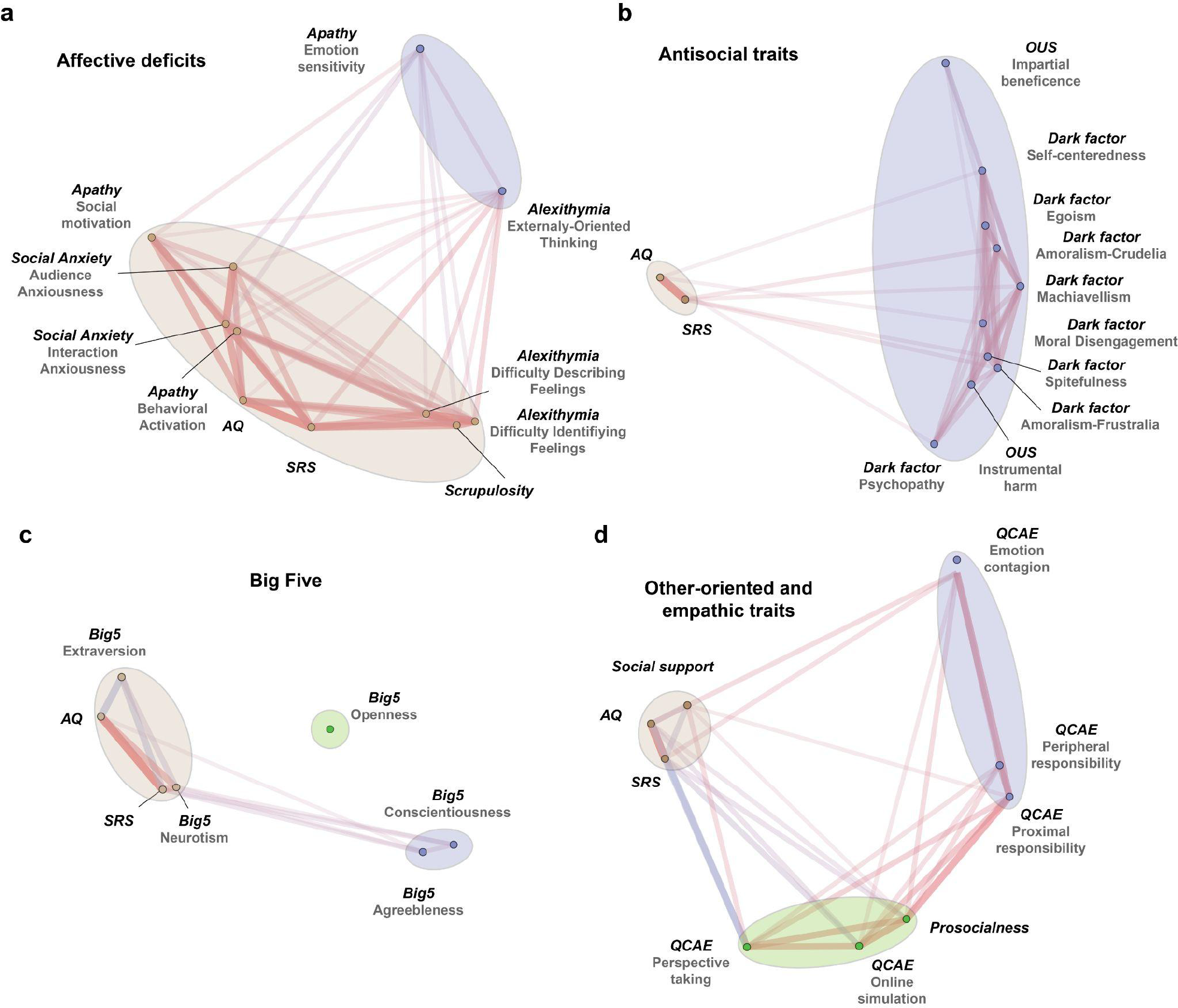
Visualization of personality trait networks. We categorized the personality measures (other than the autistic trait measures) into four groups. We then illustrated the networks consisting of autistic traits on the one hand, and each of these groups of personality measures on the other hand: **(a)** Affective deficits (including Social Anxiety, Apathy, Alexithymia, and Moral Scrupulosity), **(b)** Antisocial traits (including the Dark Factors, and utilitarianism), **(c)** the Big Five, and **(d)** Other-oriented and empathic traits (including QCAE, Perceived Social Support, and Prosocialness). Each dot in the figure indicates a personality sub-scale. Length of edges connecting the dots indicates the statistical distance (i.e., absolute correlation coefficient) between the sub-scales. Color of the edges indicates the sign of the relationship (i.e., warm color = positive association, cool color = negative association). Ellipses were drawn to reflect potential clusters in each network, which will be formally examined in the factor analysis below.

### Factor analysis

We adopted a dimension approach to personality measures and used the composite dimensional scores as a more comprehensive representation of participant personality profiles. Specifically, we carried out an exploratory factor analysis on the 33 sub-sacles from the 12 established personality questionnaires related to autistic traits, affect and social deficits, prosociality (and the lack thereof), and empathy. Using the Cattell-Nelson-Gorsuch (CNG) test, our analysis identified a 4-factor latent structure that best characterized the variance in personality data of the two groups combined (**Fig. 3a**; for details, see **Materials and Methods**). The model explained 42.1% of total variance. Based on the highest loading sub-scales (|loading| > 0.35), including the two autistic trait scores (AQ and SRS), the sub-scales of alexithymia, and social anxiety, we labeled the first factor as ‘Autistic trait and social avoidance’ (Factor 1; **Table S1**). The highest loading sub-scales for the second factor were sub-scales of trait empathy (QCAE), prosociality, and social and emotional apathy (negative loading). We therefore labeled the second factor as ‘Empathy and prosociality’ (Factor 2; **Table S2**). The third factor had the highest loading items from the Dark Triad questionnaires and was labeled as ‘Antisociality’ (Factor 3; **Table S3**). Finally, the fourth factor consisted mainly of other-regarding tendency, perceived social support, and social agreeableness, and was labeled as ‘Social agreeableness’ (Factor 4; **Table S4**).

**Figure 3.**
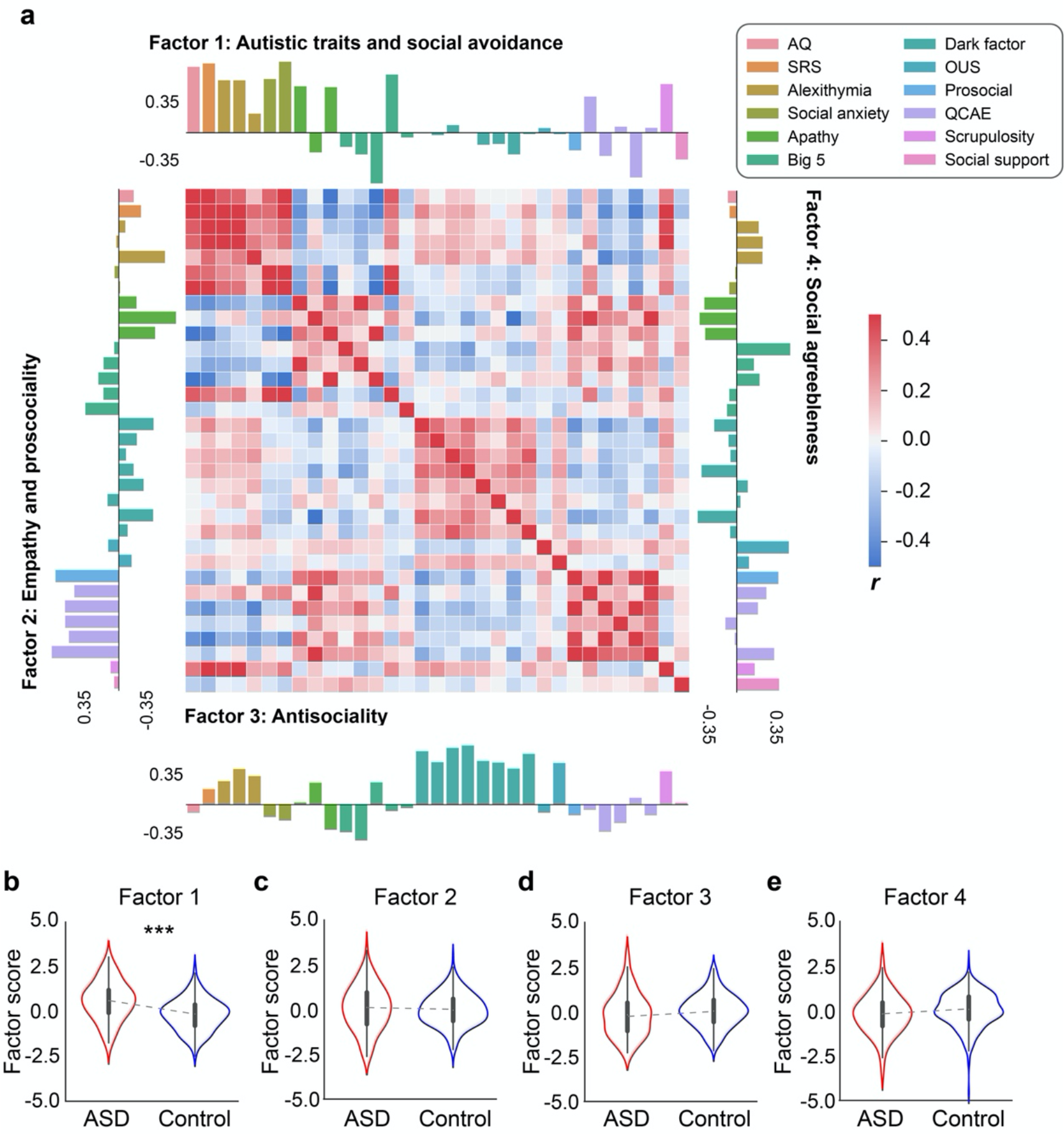
Results of factor analysis. **(a)** The correlation matrix of 33 questionnaire sub-scales and loadings of each sub-scale for the 4 factors. **(b-e)** Group differences in factor scores. The ASD group was significantly higher on Factor 1, which was primarily associated with standard autistic trait measures (i.e., AQ and SRS), social anxiety, and alexithymia. ***: *p* < 0.001

**Table 1.**
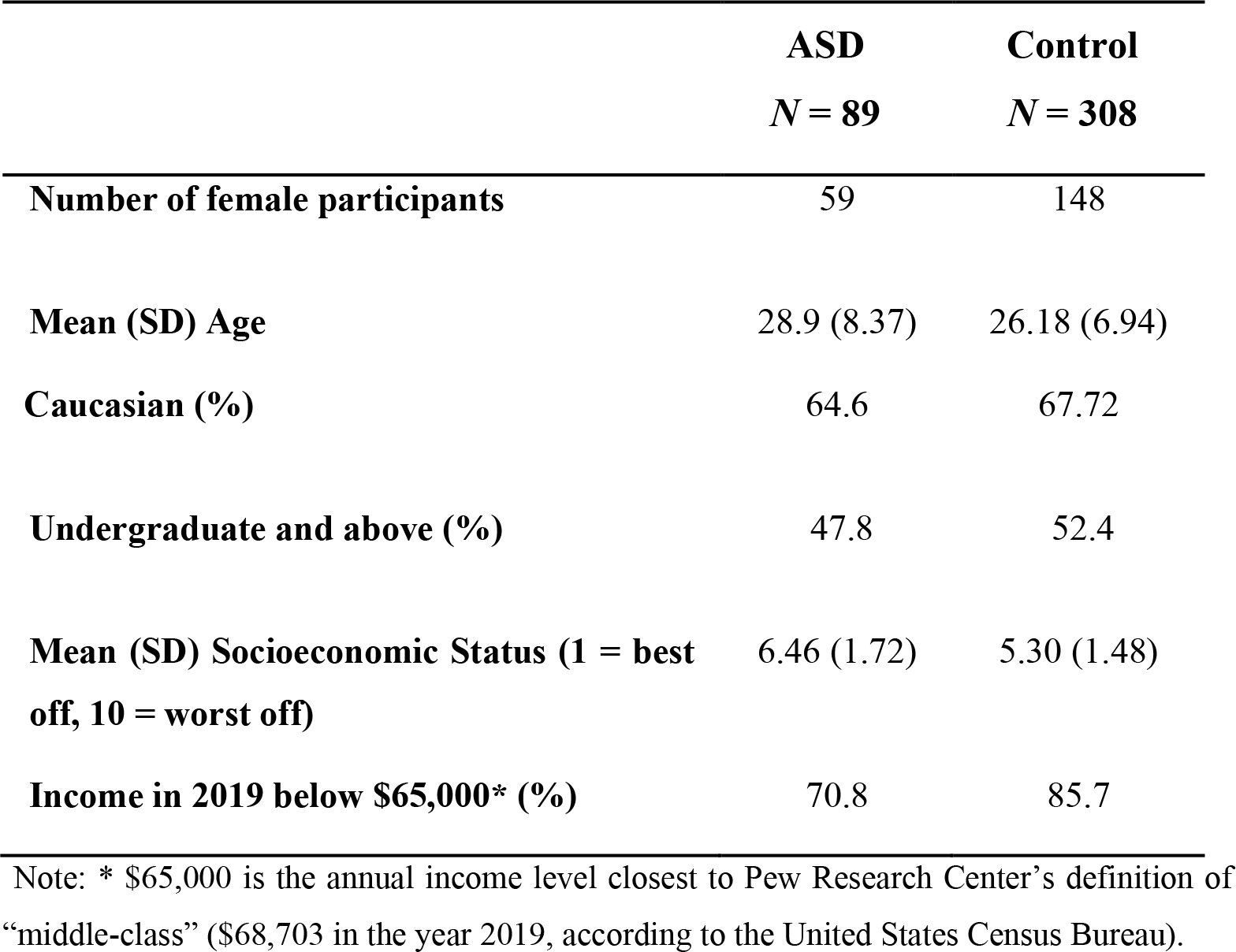
Demographics of the ASD and the control groups.

|  | **ASD**<br>*N* = 89 | **Control**<br>*N* = 308 |
| --- | --- | --- |
| **Number of female participants** | 59 | 148 |
| **Mean (SD) Age** | 28.9 (8.37) | 26.18 (6.94) |
| **Caucasian (%)** | 64.6 | 67.72 |
| **Undergraduate and above (%)** | 47.8 | 52.4 |
| **Mean (SD) Socioeconomic Status (1 = best off, 10 = worst off)** | 6.46 (1.72) | 5.30 (1.48) |
| **Income in 2019 below \$65,000\* (%)** | 70.8 | 85.7 |

Note: \* \$65,000 is the annual income level closest to Pew Research Center's definition of "middle-class" (\$68,703 in the year 2019, according to the United States Census Bureau).

**Table 2.** Correlations between factor scores and social trait judgments.

| Measure | Factor 1 |  | Factor 2 |  | Factor 3 |  | Factor 4 |  |
| --- | --- | --- | --- | --- | --- | --- | --- | --- |
|  | ASD | Control | ASD | Control | ASD | Control | ASD | Control |
| Warm | -0.12 | -0.09 | 0.15 | 0.09 | -0.23* | -0.12* | 0.02 | 0.19** |
| Critical | 0.18 | 0.07 | 0.07 | 0.04 | 0.06 | -0.10 | 0.09 | -0.03 |
| Competent | -0.20 | -0.07 | 0.13 | 0.14* | -0.24* | -0.10 | 0.06 | -0.05 |
| Practical | -0.19 | 0.03 | 0.01 | 0.10 | -0.03 | -0.00 | 0.09 | 0.11 |
| Feminine | 0.00 | -0.04 | 0.04 | 0.07 | 0.01 | -0.11 | 0.12 | -0.00 |
| Strong | -0.10 | 0.01 | 0.03 | 0.13* | 0.03 | -0.00 | 0.16 | 0.16** |
| Youthful | -0.03 | 0.00 | 0.18 | 0.16** | -0.03 | -0.15* | 0.03 | 0.06 |
| Charismatic | -0.09 | 0.00 | 0.27* | 0.18** | -0.06 | -0.12* | 0.26* | 0.15** |
| Trustworthy | -0.09 | -0.09 | 0.04 | 0.23** | -0.22* | -0.03 | 0.14 | 0.21** |
| Dominant | -0.02 | 0.02 | 0.23* | 0.17** | 0.07 | -0.00 | 0.13 | 0.21** |

Note: \* $p < 0.05$ , \*\* $p < 0.01$

To further confirm our results, we repeated the factor analysis with a randomly selected subset of ASD (41 male, 10 female) and control participants (187 male, 47 female) that approximated the gender ratio typically seen in the ASD population (i.e., male:female = 4.3:1, according to a recent report from the Centers for Disease Control and Prevention, (Maenner et al., 2020)). A qualitatively similar personality structure was obtained with this subset of participants (**Table S5** - Table S8).

In which personality dimensions do the ASD and control participants differ? To address this question, we used a linear model to compare the factor scores between groups, controlling for demographic variables including sex, age, and subjective socioeconomic status. As expected, the ASD group had significantly higher average factor scores on Factor 1 (‘Autistic trait and social avoidance’) than the control group (*B* = 0.59±0.12, *t* = 5.15, *p* < 0.001; **Fig. 3b**), demonstrating the validity of the factor analysis. All the other three factors did not exhibit any significant group difference (Factor 2: *B* = 0.06±0.12, *t* = 0.47, *p* = 0.641; Factor 3: *B* = -0.06±0.12, *t* = -0.46, *p* = 0.649; *B* = -0.19±0.13, *t* = -1.45, *p* = 0.148; **Fig. 3c-e**). It is worth noting that the ASD group did not differ from the control group on the ‘Empathy and prosociality’ dimension (i.e., Factor 2), indicating that individuals with ASD do not necessarily lack the interests in sharing others’ thoughts and feelings, or engaging in prosocial behaviors, as some influential accounts have suggested (Leekam et al., 2011; Moriuchi et al., 2017). Individuals with ASD were not higher than the control group on the ‘Antisociality’ dimension, distinguishing them from people with psychopathic and/or callous-unemotional traits (R James R Blair, 2013; Frick & Viding, 2009; Waller et al., 2020).

### No overall differences in social trait judgments across groups

Participants’ social trait judgments of faces were processed by averaging the rating of each social trait across the 50 faces that each participant saw. **Figure 1** displays the distribution of judgments along ten social traits for the ASD and the control participants. There was no significant difference in any of the mean social trait judgments between the two groups (|*t*s| < 1.63, *p*s > 0.104; **Table 2**).

### Group differences in neuronal encoding of trustworthiness and warmth

We next focused on the two social traits that are crucial for social approach tendencies, namely, trustworthiness and warmth (Fiske et al., 2007; Todorov et al., 2015). Although we did not detect differences in the average judgments of trustworthiness or warmth between the ASD group and the control group, this does not mean that the neurocognitive basis underlying the processes of these two social traits are identical for the two groups of participants. Indeed, a previous study has shown that neurons in the human amygdala and hippocampus collectively encode a social trait space, which is likely involved in the abnormal processing of social information in autism (Cao et al., 2020). Thus, to further examine the distinctions between the ASD group and the control group in terms of the underlying processing of trustworthiness and warmth, we calculated the dissimilarity matrices (DM) based on the participants’ social trait judgments, and based on the neuronal response patterns of a third group of participants, who viewed exactly the same set of faces. We then assessed the correspondence between the social trait DM and the neural response DM using the representational similarity analysis (RSA) (Kriegeskorte et al., 2008). Specifically, for this third group, we recorded from 667 neurons in the amygdala and hippocampus of 8 neurosurgical patients (23 sessions in total; overall firing rate greater than 0.15 Hz), which included 340 neurons from the amygdala, 222 from the anterior hippocampus, and 105 from the posterior hippocampus. We aligned neuronal responses at stimulus onset and used the mean normalized firing rate in a time window from 250 ms to 1250 ms after stimulus onset for subsequent analyses. Note that this third group of participants had autistic traits (measured in AQ and SRS) comparable to the control group.

We found that for trustworthiness, although the social trait DM for participants with ASD was similar to that of the control participants (**Fig. 4a**), it was less correlated with the neural response DM from the neurosurgical patients (derived from face-responsive neurons; *ρ* = -0.037 for ASD and *ρ* = 0.026 for controls; similar results were obtained when using the data from all neurons). A permutation test statistically confirmed that the difference in DM correspondence between participant groups was above chance (**Fig. 4b-d**; *p* = 0.041; see **Materials and Methods**). Similarly, although to a lesser degree, we found that for the social trait warmth the correlation between trait judgment DM and neural response DM derived from control participants was marginally significantly higher than that derived from ASD participants (**Fig. 4e-g**; *p* = 0.084). These results suggest that the processing of trustworthiness and warmth in ASD is distinguishable from that in controls at the neuronal encoding level (see **Figure S1** for the neuronal representations of the other social traits judgments).

**Figure 4.**
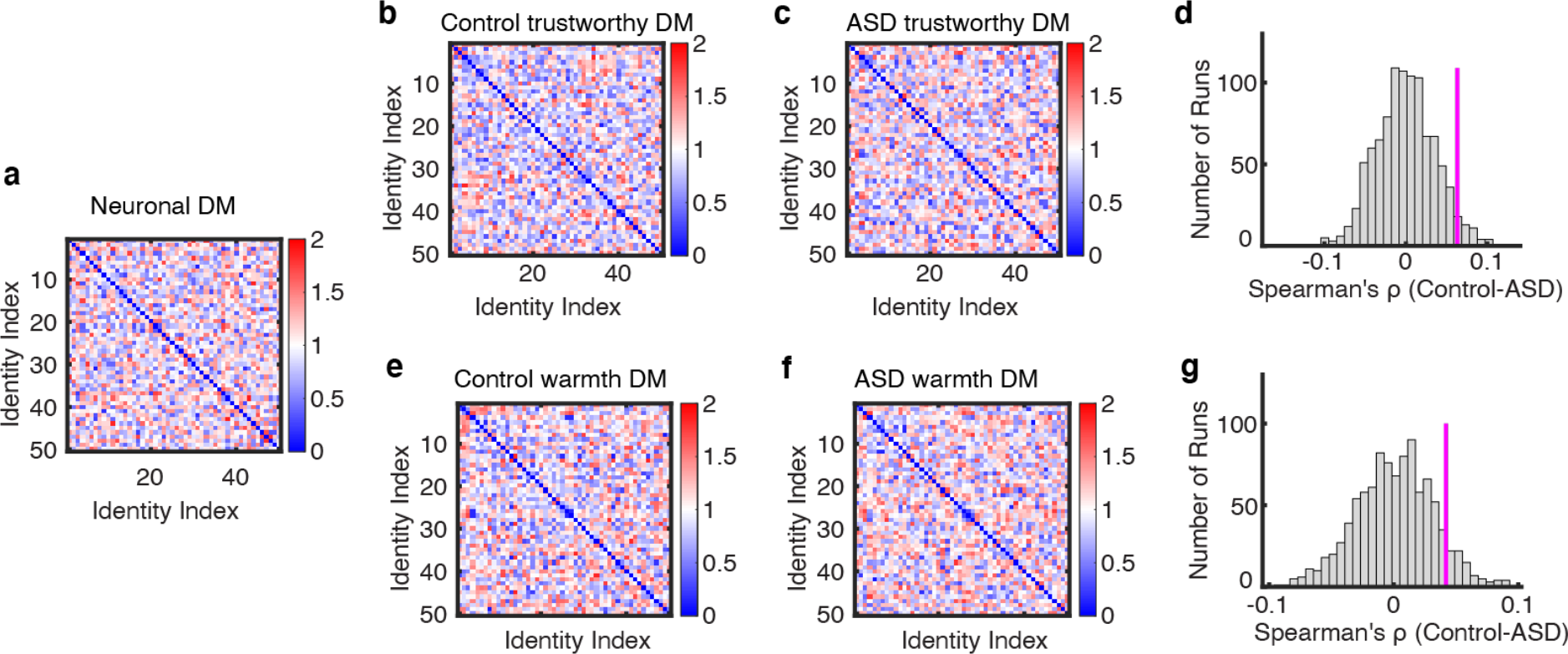
A stronger neural-rating correspondence in controls than participants with ASD. **(a)** Neuronal dissimilarity matrix (DM) constructed across face examples. **(b, c, e, f)** Social DM constructed across ratings of face examples in a single trait. **(b, e)** DM of *trustworthiness* and *warmth* from neural typical participants. **(c, f)** DM from participants with ASD. **(d, g)** Observed vs. permuted difference in DM correspondence between participant groups. The magenta line indicates the observed difference in DM correspondence between participant groups. The null distribution of difference in DM correspondence (shown in gray histogram) was calculated by permutation tests of shuffling the participant labels (1000 runs).

### Associations between personality dimensions and social trait judgments

We next examined the correlation patterns between factor scores and individual tendencies in social trait judgments for the control and the ASD participants, respectively. As illustrated in **Figure 5a** and **Table 2**, participants with ASD and controls showed qualitatively similar patterns of correlations between factor scores and social trait judgments, albeit the strengths of correlations for several traits appeared to be different between the two groups. Of note, the social avoidance and anxiety personality dimension (i.e., Factor 1), on which the two conventional autistic personality measures loaded, was not significantly correlated with any social trait judgments of faces (**Figure 5a**), a pattern that was true for both the ASD group and the control group. This suggests that social avoidance and anxiety *per se*, is unlikely to substantially contribute to the individual differences in social trait judgments from faces.

**Figure 5.**
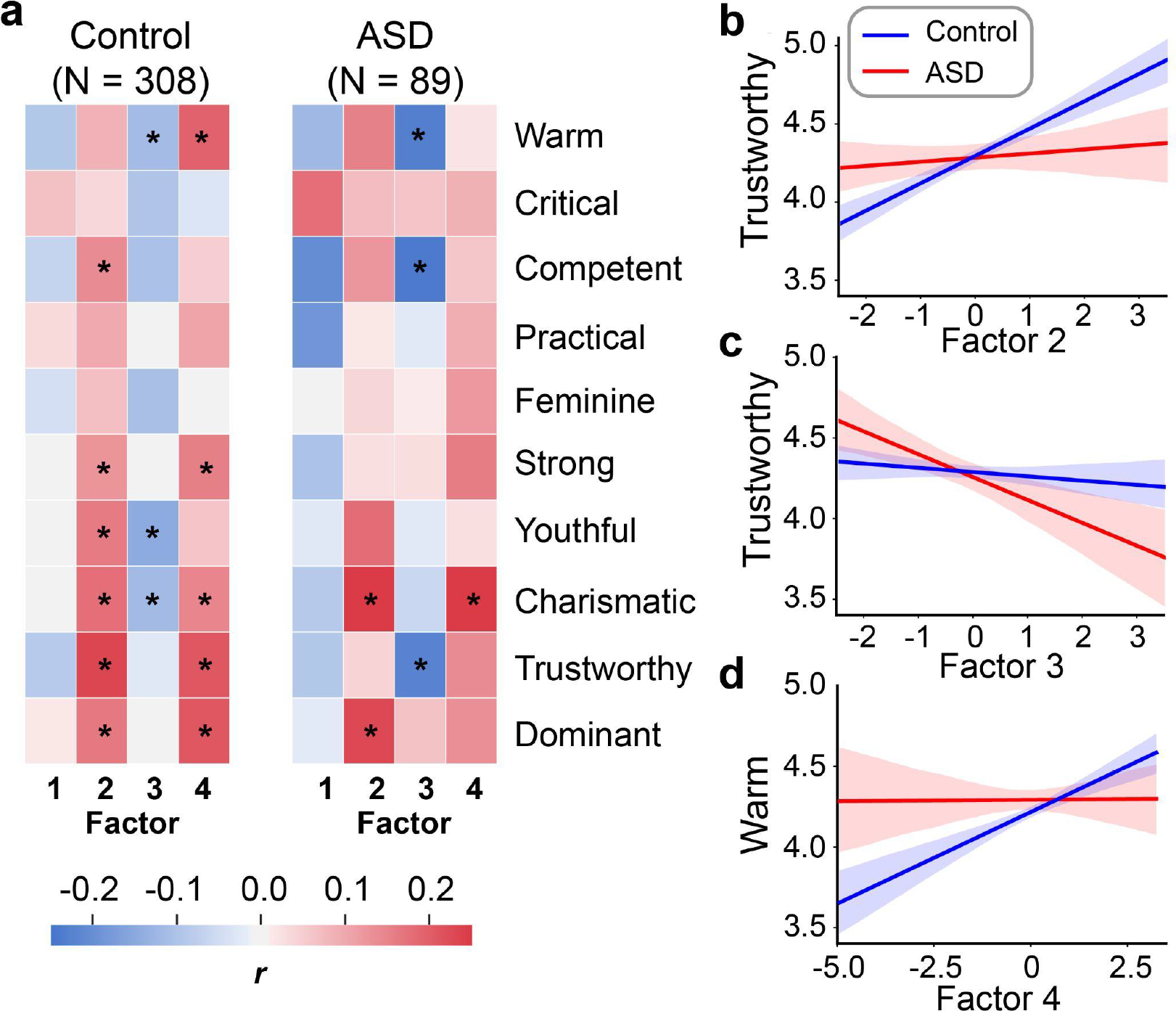
Results of the regression analysis. Scores of Factor 4 are differentially associated with the Warm judgment in the control group relative to the ASD group **(a)**. Scores of Factor 2 **(b)** and Factor 3 **(c)** are differentially associated with the Trustworthy judgment in the control group relative to the ASD group. Scores of Factor 4 (d) are differentially associated with the Warm judgment in the control group relative to the ASD group.

A comparable personality dimensional structure between groups does not necessarily imply a comparable association between personality dimensions and social trait judgments between groups. Since our neuronal encoding analysis has shown that the ASD group and the control group may have distinct neuronal representations of trustworthiness and warmth, we next examined whether the associations between personality dimensions and judgments of trustworthiness and warmth also differed between groups (**Figure 5b-d**). To this end, we ran two linear regression models to examine whether the two groups exhibited differential association patterns between personality dimensions, and trustworthiness and warmth judgments. The participants’ average ratings of trustworthiness and warmth were included as the dependent variables in the two models, respectively. The scores of all four personality dimensions (i.e., factors) were simultaneously included in the models as independent variables. Critically, we also included participant group (ASD vs. control) and the interactions between group and factor scores in the regression models. The rationale of this model was to capture the differential association between personality dimensions and trustworthiness and warmth judgments of faces between the two groups. In other words, we examined whether the associations between personality dimensions and trustworthiness and warmth judgments were attenuated or amplified in the ASD group relative to the control group. Covariates of no interest were also included (see ***Materials and Methods***).

For the regression model with trustworthiness rating, we found that the interaction between group and Factor 3 score was significant (*B*±s.e.m. = 0.15±0.08, *t* = 1.96, *p* = 0.05, CI = [-0.30, 0.02]; **Fig. 5c**). This suggests that for the participants with ASD, higher scores on the antisocial trait dimension indicates a decreased tendency to perceive a face as trustworthy (*B*±s.e.m. = -0.11±0.06, *t* = -1.76, *p* = 0.079, CI = [-0.24, 0.01]). Such an association was not observed in the control participants (*B*±s.e.m. = 0.04±0.04, *t* = 0.810, *p* = 0.418, CI = [-0.05, 0.12]). To a lesser extent, the interaction between group and Factor 2 score was marginally significant (*B*±s.e.m. = -0.13±0.07, *t* = -1.76, *p* = 0.08, CI = [-0.22, 0.09]; **Fig. 5b**). This effect indicated that the control participants who were high on the empathy and prosociality personality dimension were more likely to judge a face as trustworthy (*B*±s.e.m. = 0.13±0.05, *t* = 2.87, *p* = 0.004, CI = [0.04, 0.22]); such a relationship was absent in the participants with ASD (*B*±s.e.m. = 0.00±0.06, *t* = 0.01, *p* = 0.989, CI = [-0.12, 0.12]).

For the regression model with Warmth ratings, we found that across the two groups the main effect of Factor 3 was significantly negative (*B*±s.e.m. = -0.12±0.06, *t* = -2.16, *p* = 0.032, CI = [-0.23, - 0.01]). This suggests that participants with higher scores on the antisocial trait dimensions are less likely to judge a face as warm. Moreover, the interaction between group and Factor 4 was marginally significant (*B*±s.e.m. = 0.12±0.07, *t* = 1.83, *p* = 0.068, CI = [-0.01, 0.25]; **Fig. 5d**): control participants who had a higher social agreeableness score were more likely to perceive a face as warm (*B*±s.e.m. = 0.11±0.04, *t* = 3.17, *p* = 0.002, CI = [0.04, 0.18]); this was not the case for the participants with ASD (*B*±s.e.m. = -0.01±0.06, *t* = -0.19, *p* = 0.850, CI = [-0.12, 0.10]).

## Discussion

In this study, we combined a dimensional approach to personality, social trait judgment of faces, and neurophysiological recordings to delineate the personality dimensions underlying the distinct processing of social traits in people with ASD relative to controls. Our results suggest that the ASD and the control participants do not significantly differ along important social-affective personality dimensions (e.g., empathy, prosociality, antisociality) or social trait judgments (e.g., trustworthiness, warmth). However, two social traits that are critical to social approach tendencies, namely trustworthiness and warmth, as evaluated by the ASD and the control participants were differentially encoded neurally, and were differentially associated with social-affective personality dimensions in the ASD and the control participants. These findings contribute to the understanding of the personality profile of ASD and its relations to the atypical social trait judgments in several ways.

First, to the best of our knowledge, our study is the first to characterize the relative position of autistic traits in a comprehensive social-affective personality space. Unlike the previous research that typically investigates the relationship between autistic traits and only one or two other social-affective personalities (e.g., empathy, alexithymia), here we examined the relationship between autistic traits and ten social-affective personalities that cover empathy, prosociality, antisociality, social anxiety, and moral preferences. Given the conceptual and statistical overlap among these questionnaires, bivariate correlations would result in uninformative and problematic conclusions (cf. (Shah et al., 2019)). To address this issue, we adopted a ‘trans-diagnostic’ (dimensional) approach to personality measures (Gillan et al., 2016), applying factor analysis to the 33 sub-scales of the ten social-affective questionnaires and the two autistic trait measures (i.e., AQ and SRS). This dimensional approach allowed us to control for overlapping variance across different measurement scales, and obtain orthogonal personality dimensions. Inspecting the resultant 4-dimensional social-affective personality space, it is clear that autistic traits are most closely associated with difficulty in understanding one’s own and others’ emotions (i.e., components of alexithymia), anxiety related to social communications (i.e., social anxiety), and lack of motivation to initiate or engage in social interactions (i.e., apathy). Although previous studies have linked autistic traits with each of these social avoidance-related personality traits (Bejerot et al., 2014; Kuusikko et al., 2008; Spain et al., 2018), our results clearly demonstrated that they formed a statistically meaningful cluster, independent of other social-affective personality traits, such as empathy, prosociality, and antisociality.

With regard to empathy and prosociality, our study revealed, perhaps surprisingly, that the personality dimension characterized by autistic traits and social avoidance was statistically orthogonal to the personality dimension characterized by empathy and prosociality. We also found that participants with ASD did not differ in the personality dimension of empathy and prosociality compared to controls, which is in contrast to previous findings that empathic responding is impaired in individuals with ASD (Baron-Cohen, 2002; Dyck et al., 2001; McDonald & Messinger, 2012; White et al., 2009). One distinction needs to be made between empathic responding (probed in previous studies), which is typically measured using social interaction tasks combined with experimenter coding of empathy-related behaviors (e.g., (McDonald & Messinger, 2012)), and empathy-related attitudes and self-evaluations (probed in our current study), which are often measured by self-reported personality questionnaires. It is possible that one perceives oneself as empathetic and caring, but fails to live up to those attitudes or personal ideals in social interactions. Perhaps other personality traits, such as social anxiety and avoidance, hinder the realization of these empathetic traits. Another possibility is coder bias. Although the coders in those studies are typically blind to the purpose of the studies and unaware of the participants’ group, they are nonetheless typically developing adults. As a recent theoretical work has pointed out (Jaswal & Akhtar, 2019), the assumptions about the social meaning of certain behaviors and bodily expressions (e.g., avoiding eye-contact) that typically developing adults take for granted may not be shared by people with ASD. Such misunderstandings may result in biased evaluations of the behaviors and expressions (or the lack thereof) of people with ASD on the part of TD adult coders. Taken together, our dimensional approach reveals that autistic traits are – on the one hand – most closely related to social anxiety, avoidance, and difficulty with understanding emotions, and – on the other hand – independent of empathetic tendency, prosociality, and antisociality. Future research that combines self-reported measures, naturalistic behaviors, and the testimony of the people with ASD is needed for better ascertaining whether and in what aspects of social trait judgments people with ASD exhibit difficulties.

The second contribution of our study is to ascertain, with a reasonably powered sample, how social-affective personality dimensions are differentially related to social trait judgments of faces (e.g., warmth and trustworthiness) in people with ASD and controls. Social trait judgments of faces, in particular, warmth and trustworthiness, are crucial for social approach tendencies (Todorov et al., 2015). However, it is not clear whether and how ASD impacts social trait judgments based on the few previous studies on this topic, which typically involved very small samples (Forgeot dArc et al., 2016; Latimier et al., 2019; Lindahl, 2017). Here, combining the comprehensive social-affective personality dimensions, naturalistic face stimuli, and a set of social traits that most comprehensively characterize social judgments, we revealed that the participants with ASD exhibited altered associations between social-affective personality dimensions and social trait judgments (warmth and trustworthiness). Our finding suggests a potential psychological mechanism underlying the difficulty with social interactions observed in people with ASD, namely, self-reported empathetic and prosocial tendencies fail to translate into a way of person perception that is more conducive to social approach tendencies (i.e., perceiving others as more trustworthy and warmer). This altered social perception of other people may further hinder or discourage people with ASD to engage in social interactions. Together with a previous study (Cao et al., 2021) showing that participants with ASD have altered neuronal representations of social traits in the amygdala and hippocampus (replicated in this study), we speculate that the distinct neural encoding of social traits may lead to a different set of criteria (or thresholds) for judging a face as trustworthy and warm by people with ASD.

One limitation of our study is the sample size of the ASD group (*N* = 89), although we note that this sample size was more than three times of most of the previous studies investigating the social trait judgments from faces in people with ASD (average *N* = 28; (Forgeot dArc et al., 2016; Latimier et al., 2019; Lindahl, 2017)). Moreover, we found that the Factor 3 score (antisocial traits) was significantly more (negatively) correlated with less trustworthiness judgments in the ASD group than in the control group (**Fig. 4c**), despite the ASD group having fewer participants. This complements the above speculation, that ASD may have a distinct criteria/threshold for judging trustworthiness and warmth, perhaps due to their differences in underlying neural processing. Future research with better powered ASD samples, perhaps via multi-center collaborations (Di Martino et al., 2017; Valk et al., 2015), is needed to replicate and extend our findings. Lastly, it is worth noting that our participants with ASD are self-identified and we were not able to verify their autism diagnoses. However, their behavior and autism demonstration are in line with a small sample of ASD participants from our laboratory who have confirmed diagnoses. Our approach combining online recruitment / crowdsourcing with in-lab testing will be valuable to study psychiatric symptoms in social, behavioral, and clinical sciences (Shapiro et al., 2013). A future study is needed to further replicate our present findings in a large sample of ASD participants with confirmed diagnoses.

In conclusion, by integrating neuronal recording, social trait judgments, and recent advances in the dimensional approach to personality, we characterized a comprehensive social-affective personality space and ascertained the relative position of autistic traits in this space. We found that autistic traits were most closely associated with social anxiety, avoidance, and difficulty with understanding emotions, but were orthogonal to empathetic traits, prosociality, antisociality, and moral preferences. These novel personality trait dimensions further revealed altered patterns of individual differences in the judgments of trustworthiness and warmth of faces in people with ASD compared with controls, thereby shedding new light on the psychological mechanisms underlying the difficulties with social interactions and communications central to ASD.

## Acknowledgements

We thank all participants for their participation, staff from WVU Ruby Memorial Hospital for support with patient testing, and Paula Webster for participant recruitment and valuable comments. We thank Dr. Muyu Lin for her helpful comments on statistical analysis of the personality data, and Mr. Nicholas Kim for proofreading an earlier version of the manuscript. This research was supported by an NSF CAREER Award (BCS-1945230), Air Force Young Investigator Program Award (FA9550-21-1-0088), Dana Foundation Clinical Neuroscience Award, and ORAU Ralph E. Powe Junior Faculty Enhancement Award (to S.W.).

## Supplementary Materials

### Supplementary Methods

#### Single-neuron recordings and neuronal response to faces

We recorded from implanted depth electrodes in the amygdala and hippocampus from patients with pharmacologically intractable epilepsy. Target locations in the amygdala and hippocampus were verified using post-implantation CT. At each site, we recorded from eight 40 μm microwires inserted into a clinical electrode as described previously (Rutishauser et al., 2013; Rutishauser, Mamelak, et al., 2006). Efforts were always made to avoid passing the electrode through a sulcus, and its attendant sulcal blood vessels, and thus the location varied but was always well within the body of the targeted area. Microwires projected medially out at the end of the depth electrode and examination of the microwires after removal suggests a spread of about 20-30 degrees. The amygdala electrodes were likely sampling neurons in the mid-medial part of the amygdala and the most likely microwire location is the basomedial nucleus or possibly the deepest part of the basolateral nucleus. Bipolar wide-band recordings (0.1-9000 Hz), using one of the eight microwires as the reference, were sampled at 32 kHz and stored continuously for off-line analysis with a Neuralynx system. The raw signal was filtered with a zero-phase lag 300-3000 Hz bandpass filter and spikes were carefully sorted using a semi-automatic template matching algorithm as described previously (Rutishauser, Schuman, et al., 2006).

We used a 1-back task to acquire neural responses to the same CelebA stimuli from neurosurgical patients. In each trial, a single face was presented at the center of the screen for a fixed duration of 1 second, with uniformly jittered inter-stimulus-interval (ISI) of 0.5-0.75 seconds. Each image subtended a visual angle of approximately 10°. A simple 1-back task required patients to press a button if the present face image was *identical* to the immediately previous image. Nine percent of the trials were one-back repetitions. Each face was shown once unless repeated in one-back trials; and the faces shown in one-back trials were randomly selected for each patient. We excluded responses from one-back trials to have an equal number of responses for each face. This task kept patients attending to the faces, but avoided potential biases from focusing on any particular facial feature (e.g., the color of their eyes or whether they are smiling) or social judgment (e.g., whether they seem happy). The order of faces was randomized for each patient. Stimuli were presented resolution: 1600 × 1280).

Only units with an average firing rate of at least 0.15 Hz during the entire task were considered. Only single units were considered. Trials were aligned to stimulus onset. We used the mean firing rate in a time window 250 ms to 1250 ms after stimulus onset as the response to each face. Firing rate was then normalized by dividing the mean activity in the baseline (−250 ms to 0 ms relative to stimulus onset). Such normalization was applied in previous studies that analyzed the similarity between single-neuron responses to visual categories (Reber et al., 2019).

#### Supplementary Results

For completeness, here we reported the results of the analysis of the neurophysiological data with regard to all 10 social traits. We constructed a social dissimilarity matrix (DM) between identity pairs by computing covariance on the traits between each pair of face identities for control (Fig. S1a) and ASD group (Fig. S1b), and the neural DM by calculating the covariance across neurons (Fig. S1c). We used a bootstrap with 1000 runs to estimate the distribution of DM correspondence for each participant group. In each run, 70% of the data were randomly selected from each participant group and we calculated the correspondence (Spearman’s ρ) between the social trait DM and the neural response DM for each participant group. We then created a distribution of DM correspondence for each participant group. Similar to the analysis of single traits above, we used a permutation test to determine whether there was a significant difference between the correspondence of neural and social DMs in participants with ASD and controls.

We found that the observed correspondence difference between the ASD and control groups is significantly larger than when the difference was estimated with label-shuffled data (permutation p < 0.001). In addition, we also used a bootstrapping approach to estimate the distribution of DM correspondence for each participant group and we found that the two distributions were largely separated (Fig. S1d; the mean of the ASD distribution was significantly outside the control distribution [p < 0.011] and the mean of the control distribution was also significantly outside the ASD distribution [p < 0.017]).

### Supplementary Figures

**Figure S1.**
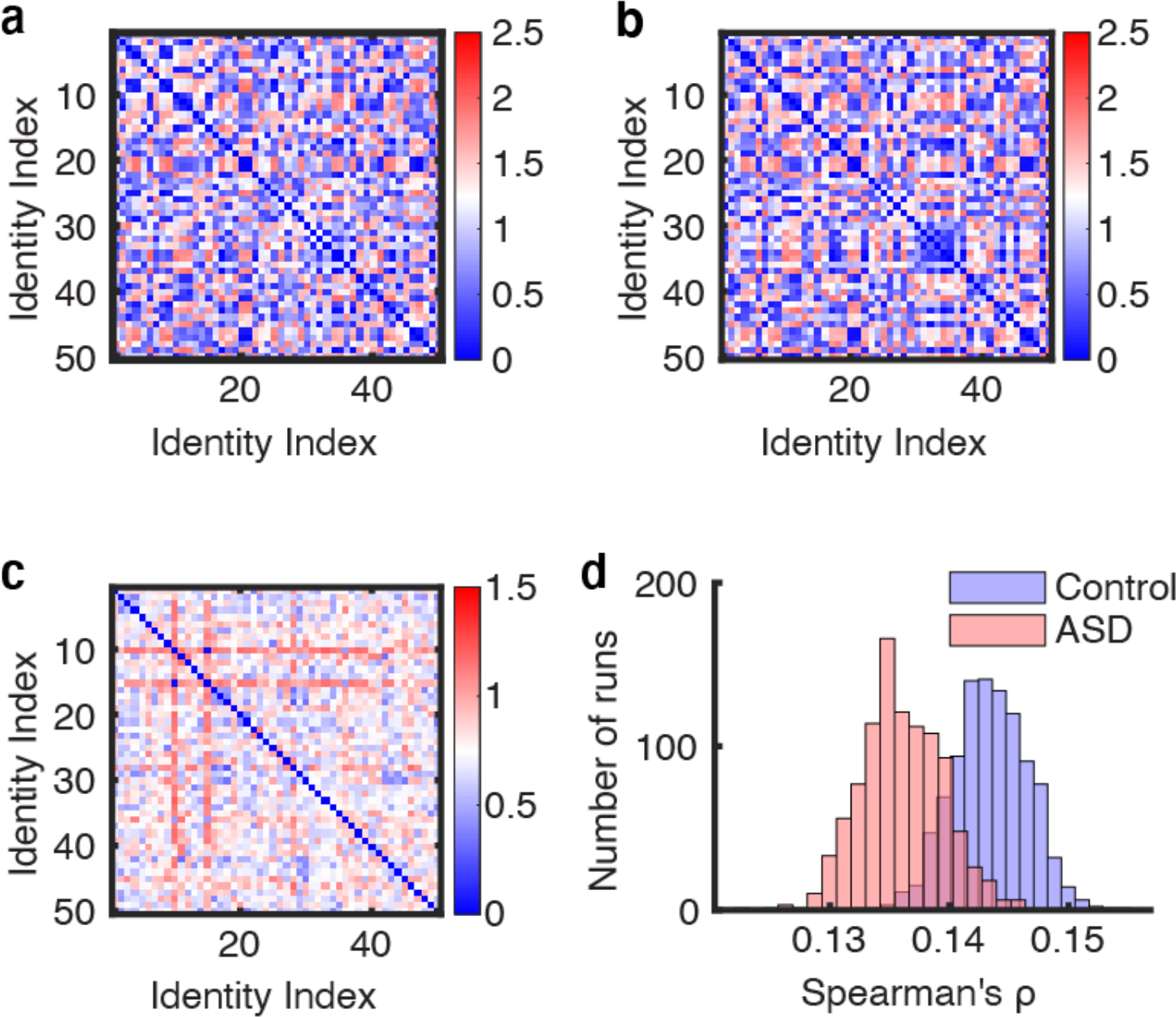
Results of the analysis of the neurophysiological data with regard to all ten social traits. **(a, b)** Average social DM constructed across 10 social traits for control and ASD using consensus ratings acquired from each run of the bootstrapping. **(c)** Neuronal DM constructed across neurons using the average response of each identity. **(d)** Bootstrap distribution of DM correspondence for each participant group. Blue: online controls. Red: online participants self-identified as ASD. Participants with ASD showed a weaker correspondence with the neural response DM compared to controls.

**Figure S2.**
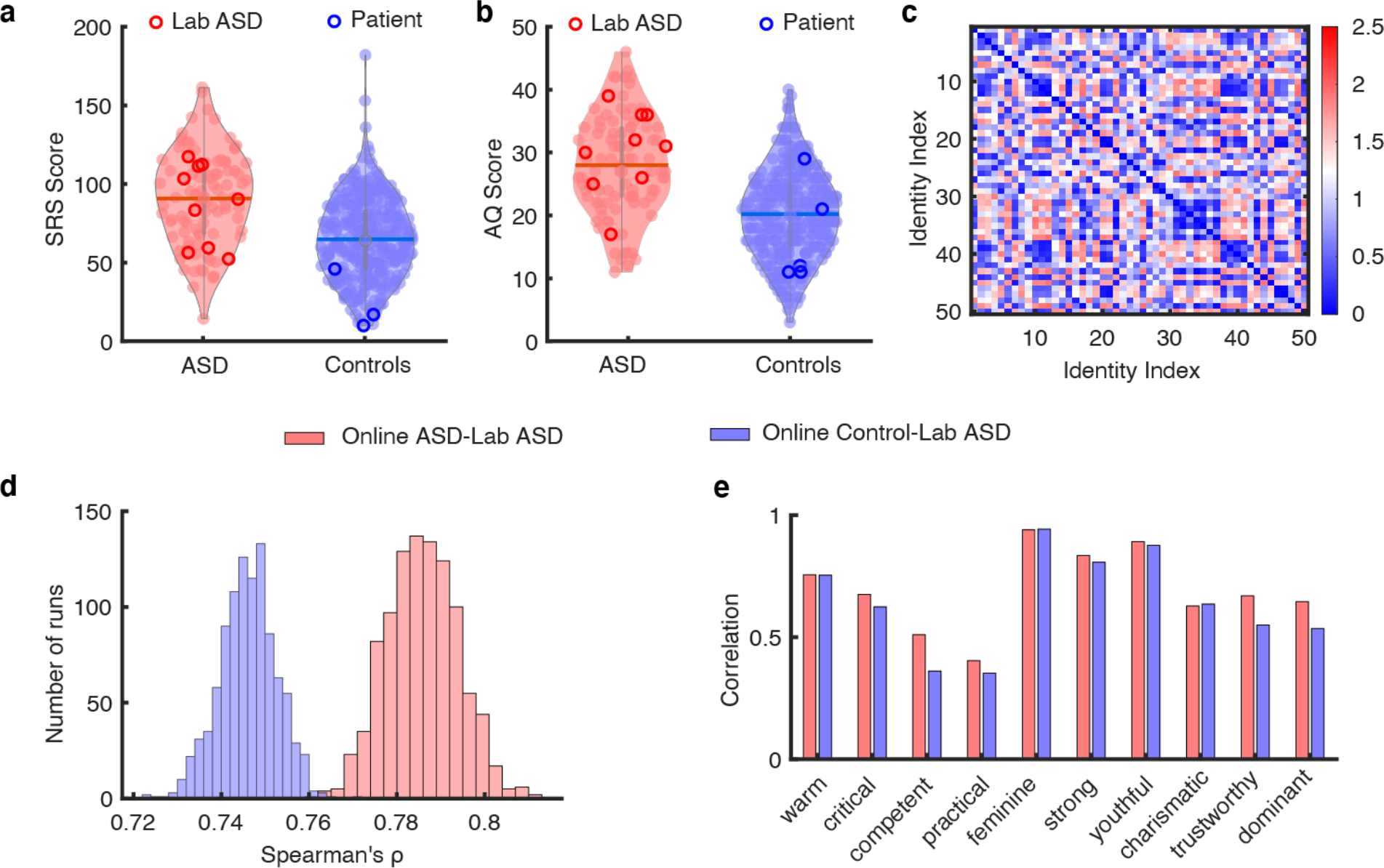
Comparable autristic scores and face judgements between participants with crowd-source ASD and in-lab participants with diagnosed ASD. **(a, b)** The violin plots illustrate the distribution of SRS scores **(a)** and AQ scores **(b)** for online participants self-identified with ASD and online control participants. The dark red circles indicate the SRS and AQ scores from in-lab participants with diagnosed ASD. The dark blue circles indicate scores of intracranial (neurotypical) patients. **(c)** Social trait DM constructed across 10 traits of ratings from in-lab participants with ASD. The social trait DM shows dissimilarity between all pairs of face identities and reflects an overall structure of the social trait judgment space. **(d)** Bootstrap distribution of DM correspondence between in-lab participants with crowd-source participants with ASD (red) and controls (blue). **(e)** Face judgements are highly correlated between in-lab participants with ASD (red) and online crowd-source participants with ASD (blue).

#### Supplementary Tables

**Table S1.** Top loading items on Factor 1 (‘Autistic traits and social avoidance’): full sample.

| Questionnaire subscale | Loading |
| --- | --- |
| Social Anxiety: Interaction anxiousness | 0.842 |
| SRS | 0.822 |
| AQ | 0.782 |
| Big 5: Neuroticism | 0.690 |
| Social Anxiety: Audience anxiousness | 0.637 |
| Alexithymia: Difficulty describing emotion | 0.622 |
| Alexithymia: Difficulty identifying emotion | 0.621 |
| Moral Scrupulosity | 0.577 |
| QCAE: Emotion contagion | 0.429 |
| Big 5: Extraversion | 0.601 |
| Apathy: Behavioural motivation | 0.551 |
| Apathy: Social motivation | 0.540 |
| QCAE: Perspective taking | 0.531 |

Note: Summary of item loadings onto the 'Autistic traits and social avoidance' factor. The top loading subscales from each questionnaire ( $|\text{loading}| > 0.35$ ) are displayed in descending order. Items with red loadings numbers are reverse-coded. SRS = Social Responsiveness Scale, AQ = Autism-spectrum Quotient (AQ), QCAE = Questionnaire of Cognitive and Affective Empathy.

**Table S2.** Top loading items on Factor 2 (‘Empathy and Prosociality’): full sample.

| Questionnaire subscale | Loading |
| --- | --- |
| QCAE: Proximal responsivity | 0.743 |
| Prosocialness | 0.702 |
| Apathy: Emotional sensitivity | 0.633 |
| QCAE: Online simulation | 0.596 |
| QCAE: Peripheral responsivity | 0.594 |
| QCAE: Perspective taking | 0.553 |
| QCAE: Emotion contagion | 0.493 |
| Apathy: Social motivation | 0.402 |
| Big 5: Openness | 0.371 |
| Alexithymia: Externally oriented | 0.512 |
| Dark Factor: Crudelia | 0.551 |
| Dark Factor: Self-centeredness | 0.385 |
| Dark Factor: Moral Disengagement | 0.380 |

Note: Summary of item loadings onto the 'Empathy and Prosociality' factor. The top loading subscales from each questionnaire ( $|loading| > 0.35$ ) are displayed in descending order. Items with red loadings numbers are reverse-coded. QCAE = Questionnaire of Cognitive and Affective Empathy.

**Table S3.**
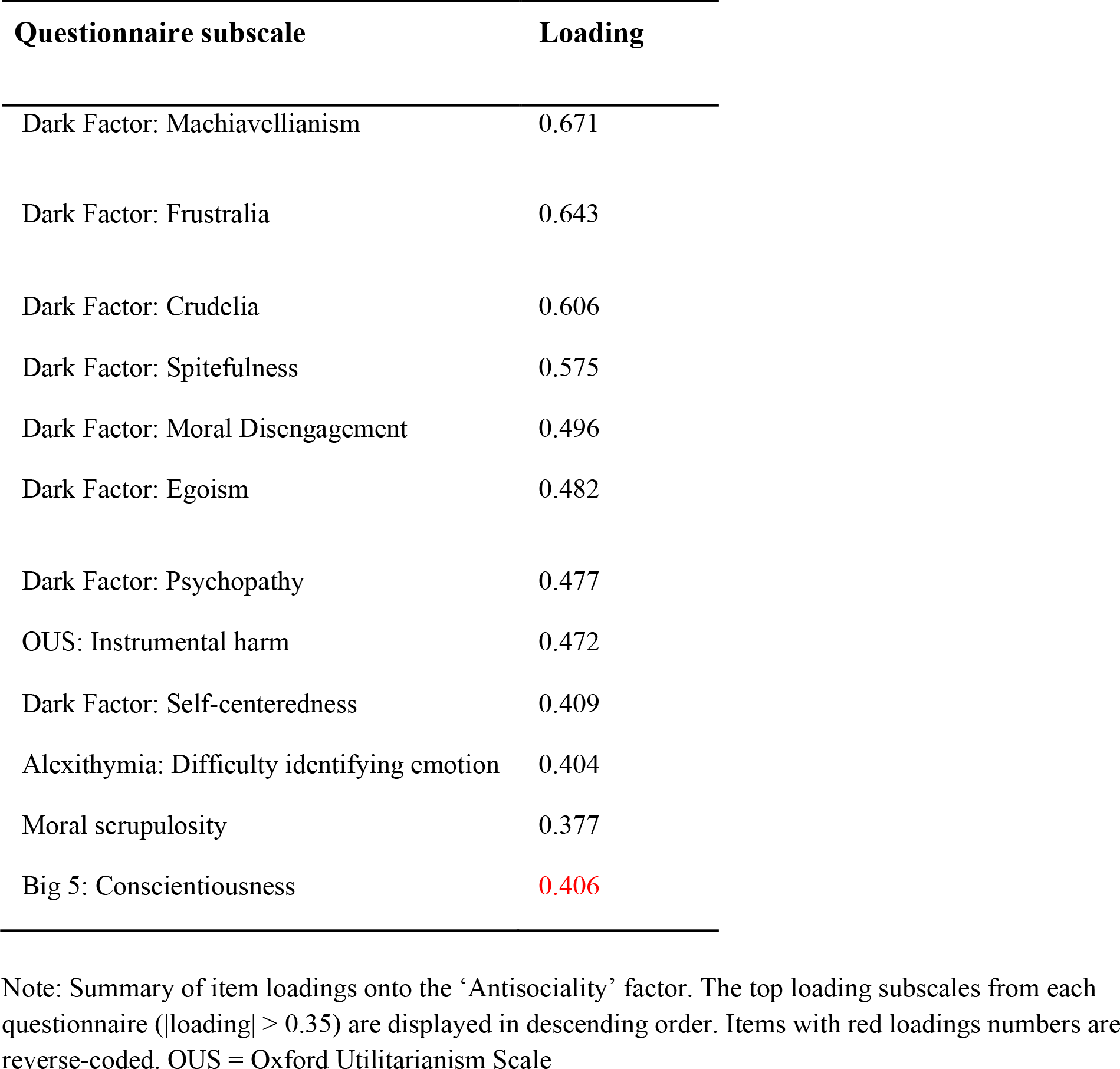
Top loading items on Factor 3 (‘Antisociality’): full sample.

**Table S4.** Top loading items on Factor 4 (‘Social agreeableness’): full sample.

| **Questionnaire subscale** | **Loading** |
| --- | --- |
| Big 5: Agreeableness | 0.552 |
| OUS: Impartial beneficence | 0.537 |
| Social support | 0.436 |
| Prosocialness | 0.429 |
| QCAE: Proximal responsivity | 0.386 |
| Apathy: Emotional sensitivity | 0.383 |
| Dark Factor: Self-centeredness | 0.404 |
| Dark Factor: Machiavellianism | 0.363 |

Note: Summary of item loadings onto the ‘Social agreeableness’ factor. The top loading subscales from each questionnaire ( $|\text{loading}| > 0.35$ ) are displayed in descending order. Items with red loadings numbers are reverse-coded. OUS = Oxford Utilitarianism Scale, QCAE = Questionnaire of Cognitive and Affective Empathy.

**Table S5.** Top loading items on Factor 1 (‘Autistic traits and social avoidance’): sex ratio matched sample.

| Questionnaire subscale | Loading |
| --- | --- |
| Social Anxiety: Interaction anxiousness | 0.814 |
| SRS | 0.710 |
| AQ | 0.691 |
| Big 5: Neuroticism | 0.671 |
| Social Anxiety: Audience anxiousness | 0.620 |
| Alexithymia: Difficulty describing emotion | 0.618 |
| Alexithymia: Difficulty identifying emotion | 0.599 |
| Moral Scrupulosity | 0.599 |
| QCAE: Emotion contagion | 0.589 |
| Apathy: Emotional sensitivity | 0.417 |
| Big 5: Extraversion | 0.485 |
| Apathy: Behavioral activation | 0.409 |
| Dark Factor: Self-centeredness | 0.408 |

Note: Summary of item loadings onto the 'Autistic traits and social avoidance' factor. The top loading subscales from each questionnaire ( $|\text{loading}| > 0.35$ ) are displayed in descending order. Items with red loadings numbers are reverse-coded. SRS = Social Responsiveness Scale, AQ = Autism-spectrum Quotient (AQ), QCAE = Questionnaire of Cognitive and Affective Empathy.

**Table S6.** Top loading items on Factor 2 (‘Empathy and Prosociality’): sex ratio matched sample.

| Questionnaire subscale | Loading |
| --- | --- |
| QCAE: Proximal responsivity | 0.666 |
| QCAE: Perspective taking | 0.621 |
| Prosocialness | 0.615 |
| QCAE: Peripheral responsivity | 0.611 |
| Apathy: Emotional sensitivity | 0.545 |
| Apathy: Social motivation | 0.525 |
| QCAE: Online simulation | 0.502 |
| QCAE: Emotion contagion | 0.429 |
| Big 5: Openness | 0.393 |
| Big 5: Extraversion | 0.369 |
| Alexithymia: Externally oriented | 0.447 |

Note: Summary of item loadings onto the 'Empathy and Prosociality' factor. The top loading subscales from each questionnaire ( $|\text{loading}| > 0.35$ ) are displayed in descending order. Items with red loadings numbers are reverse-coded. QCAE = Questionnaire of Cognitive and Affective Empathy.

**Table S7.**
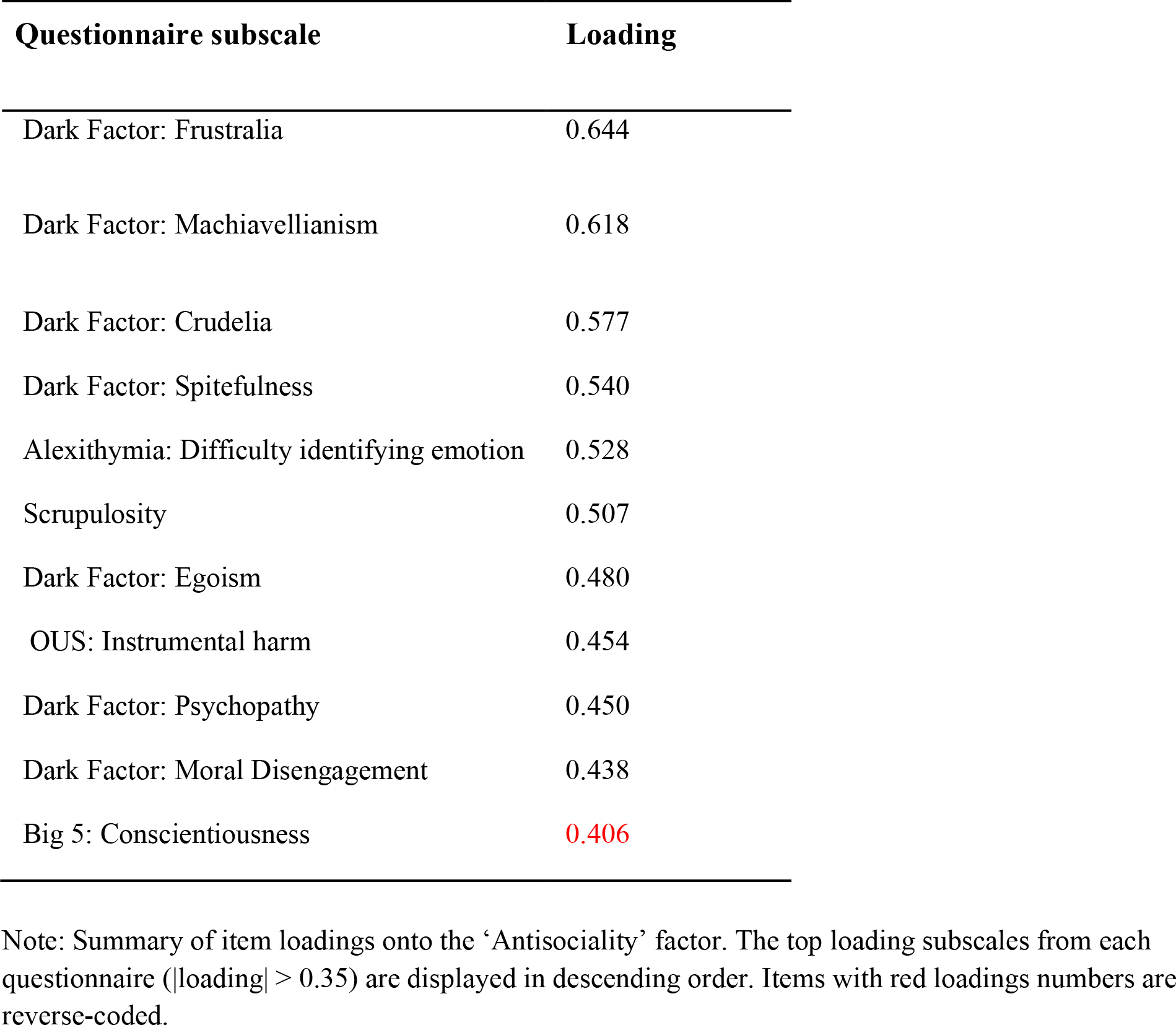
Top loading items on Factor 3 (‘Antisociality’): sex ratio matched sample.

**Table S8.** Top loading items on Factor 4 (‘Social agreeableness’): sex ratio matched sample.

| Questionnaire subscale | Loading |
| --- | --- |
| Big 5: Agreeableness | 0.567 |
| Apathy: Behavioural activation | 0.542 |
| OUS: Impartial beneficence | 0.461 |
| Social support | 0.412 |
| Big 5: Conscientiousness | 0.389 |
| Prosocialness | 0.381 |

Note: Summary of item loadings onto the ‘Social agreeableness factor. The top loading subscales from each questionnaire ( $|\text{loading}| > 0.35$ ) are displayed in descending order. OUS = Oxford Utilitarianism Scale.

